# Non-integrating direct reprogramming generates therapeutic endothelial cells with sustained vascular regeneration capacity enhanced by nanomatrix delivery

**DOI:** 10.1101/2025.09.24.678308

**Authors:** Sangho Lee, Dandan Chen, Yoshiaki Tanaka, Seongho Bae, Han Jacob Li, Er Yearn Jang, Eric Chan, In-Hyun Park, Ho-Wook Jun, Changwon Park, Young-sup Yoon

## Abstract

**Rationale:** Direct reprogramming of fibroblasts into endothelial cells (rECs) using ETV2 shows promise for vascular regeneration. However, current approaches using integrating viral vectors pose clinical translation barriers, and poor long-term cell survival limits therapeutic efficacy.

**Objective:** To develop a clinically compatible method for generating rECs using non-integrating adenoviral ETV2 (Ad-ETV2) and enhance their engraftment and therapeutic efficacy through peptide amphiphile (PA) nanomatrix encapsulation.

**Methods and Results:** Human dermal fibroblasts were reprogrammed using Ad-ETV2 and characterized by flow cytometry, RNA sequencing, and functional assays. Therapeutic efficacy was evaluated in murine hindlimb ischemia with or without PA-RGDS encapsulation over 12 months. Ad-ETV2 induced robust endothelial gene expression (CDH5, KDR, PECAM1) within 6 days, with 40-50% reprogramming efficiency. KDR+ Ad-rECs demonstrated functional endothelial properties including Ac-LDL uptake, tube formation, and exceptional proangiogenic factor secretion (200-fold higher HGF than HUVECs). RNA sequencing revealed rapid transcriptional reprogramming with fibroblast gene suppression and endothelial/angiogenic gene activation. In hindlimb ischemia, Ad-rECs significantly enhanced blood flow recovery and capillary density versus controls. Long-term analysis revealed sustained vascular contribution through three mechanisms: direct incorporation, perivascular support, and vessel guidance, persisting throughout 12 months—the longest reported follow-up for reprogrammed cells. PA-RGDS encapsulation markedly improved cell retention; while 75% of cells were lost by 3 months, retention stabilized thereafter with minimal additional loss through 12 months.

**Conclusions:** Adenoviral ETV2 delivery enables efficient generation of clinically compatible rECs without genomic integration. These cells demonstrate potent and sustained therapeutic efficacy through multiple vascular regeneration mechanisms. PA-RGDS encapsulation significantly enhances long-term engraftment, establishing this combined approach as a promising platform for treating ischemic cardiovascular diseases.

## INTRODUCTION

Cardiovascular diseases (CVDs) remain the leading cause of morbidity and mortality in industrialized nations^1,2^. Despite advances in therapeutic interventions, treatment options for patients with severe ischemic CVDs remain limited. The primary pathophysiology underlying ischemic CVD involves vascular occlusion and consequent reduced blood perfusion. Consequently, therapeutic neovascularization inducing new blood vessels has emerged as a promising strategy^3,4^.

Human induced pluripotent stem cell-derived endothelial cells (hiPSC-ECs) have demonstrated therapeutic neovascularization potential. However, clinical translation faces significant barriers including tumorigenicity risks, potential for aberrant tissue formation, complex differentiation protocols, and low efficiency^5–12^. Direct cellular reprogramming offers an alternative strategy for generating target cells in vivo and in vitro. By employing lineage-specific transcription factors, this approach bypasses pluripotency, potentially reducing adverse effects while streamlining target cell generation^6,13^. Recently, we and others have successfully generated endothelial cells through direct reprogramming using endothelial-specific transcription factors^14–17^.

ETV2, an ETS family transcription factor, plays essential roles in vascular endothelial specification and development during early murine embryogenesis^18–25^. We previously demonstrated that ETV2 alone could directly reprogram postnatal human dermal fibroblasts (HDFs) into functional endothelial cells (rECs)^16^. Upon ETV2 transduction, HDFs underwent morphological transformation and acquired endothelial gene and protein expression patterns^16^. These rECs exhibited characteristic endothelial properties *in vitro* and demonstrated vessel-forming capacity with therapeutic efficacy in ischemic animal models. While other groups have shown that various transcription factor combinations including ETV2 can reprogram human somatic cells into EC-like cells with neovascularization potential^15–17,26,27^, our previous study uniquely demonstrated that ETV2 alone under specific culture conditions could generate functional rECs capable of engrafting for three months *in vivo*—the longest reported survival among directly reprogrammed endothelial cells^16^. Critically, all existing studies, including ours, relied on lentiviral or retroviral vectors for transcription factor delivery. These integrating vectors pose significant clinical translation barriers due to permanent genomic integration of exogenous sequences, raising concerns about insertional mutagenesis and long-term safety. Clinical application of rEC therapy therefore requires development and validation of non-integrating delivery systems that maintain robust reprogramming efficiency while eliminating genomic modification risks.

For clinical translation, poor cell survival represents a critical barrier to effective cell therapy^28–32^. Studies consistently demonstrate that transplanted cells largely disappear within one week, substantially limiting therapeutic efficacy^33–36^. The hostile inflammatory environment and insufficient extracellular matrix support contribute to rapid cell death or washout. Biomaterial-based delivery systems have shown promise in enhancing cell engraftment. We developed an extracellular matrix-mimicking peptide amphiphile nanomatrix (PA-RGDS) gel that enhances survival of encapsulated cells, including pluripotent stem cell-derived cardiomyocytes and endothelial cells^37,38^. However, biomaterial-mediated delivery of directly reprogrammed ECs has not been investigated in cardiovascular disease models.

Here, we employed adenoviral vectors to deliver ETV2 for reprogramming HDFs into endothelial cells. These adenovirally-reprogrammed ECs (Ad-rECs) exhibited comparable characteristics to lentivirally-reprogrammed cells^16^ while offering better clinical compatibility. Ad-rECs demonstrated robust neovascularization capacity and therapeutic efficacy in ischemic models. Furthermore, PA-RGDS encapsulation significantly enhanced long-term cell engraftment, survival, and neovascularization over twelve months—the longest follow-up reported for directly reprogrammed cells.

## RESULTS

### Generation of directly reprogrammed endothelial cells using adenoviral ETV2

To develop clinically-compatible rECs, we constructed an adenoviral vector harboring human ETV2 (Ad-ETV2) using the Ad5 system (Supplementary Fig. S1). Defective type 5 adenovirus vectors have been widely used in clinical trials and were shown to be safe^39^. HDFs were transduced at MOI 70, optimized for maximal EC gene induction with minimal cytotoxicity, then cultured in reprogramming medium supplemented with ascorbic acid (Vit.C) and a TGFβ signaling pathway inhibitor, SB 431542 (SB) (Fig. 1A).

**Figure 1.**
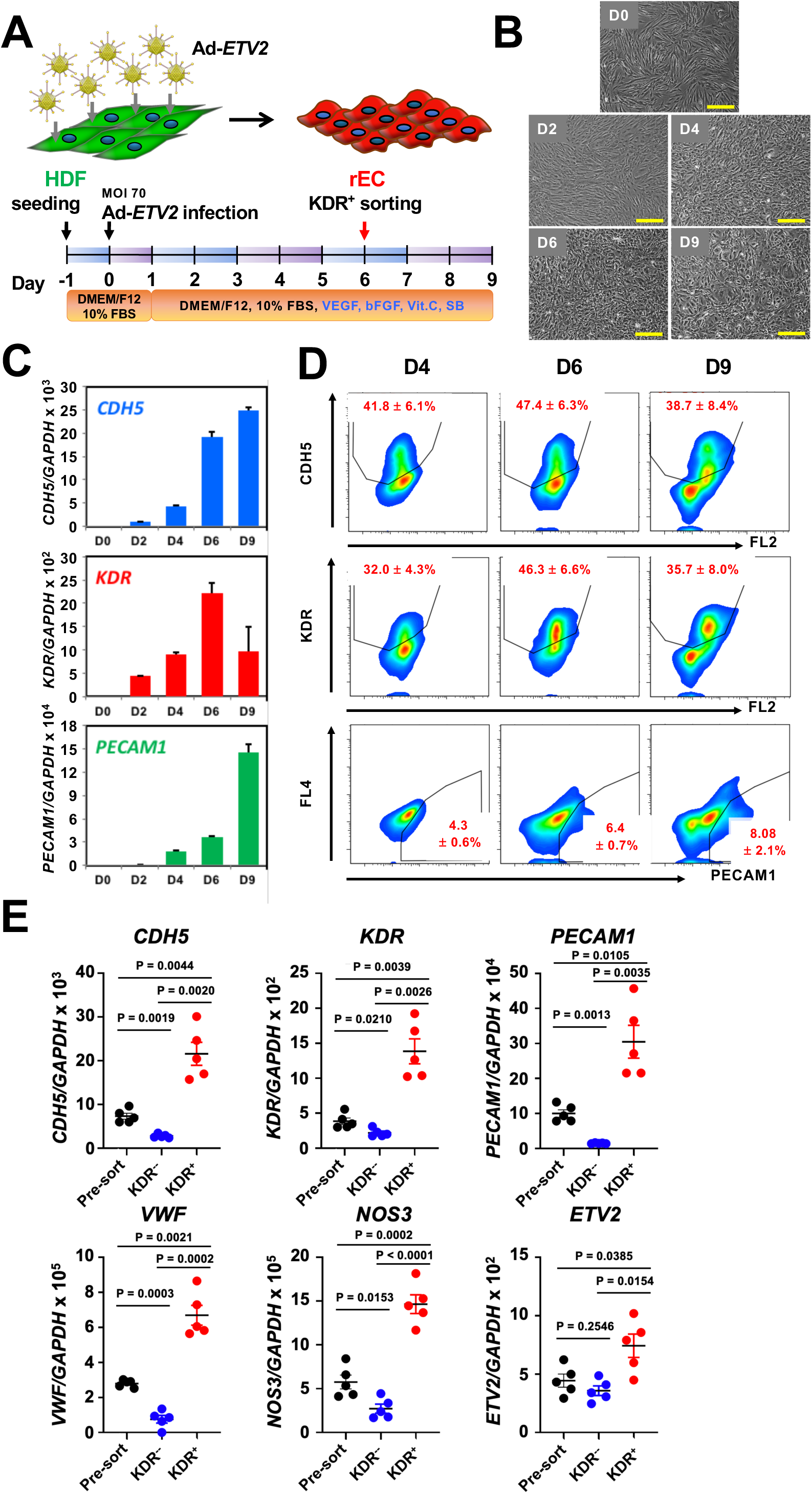
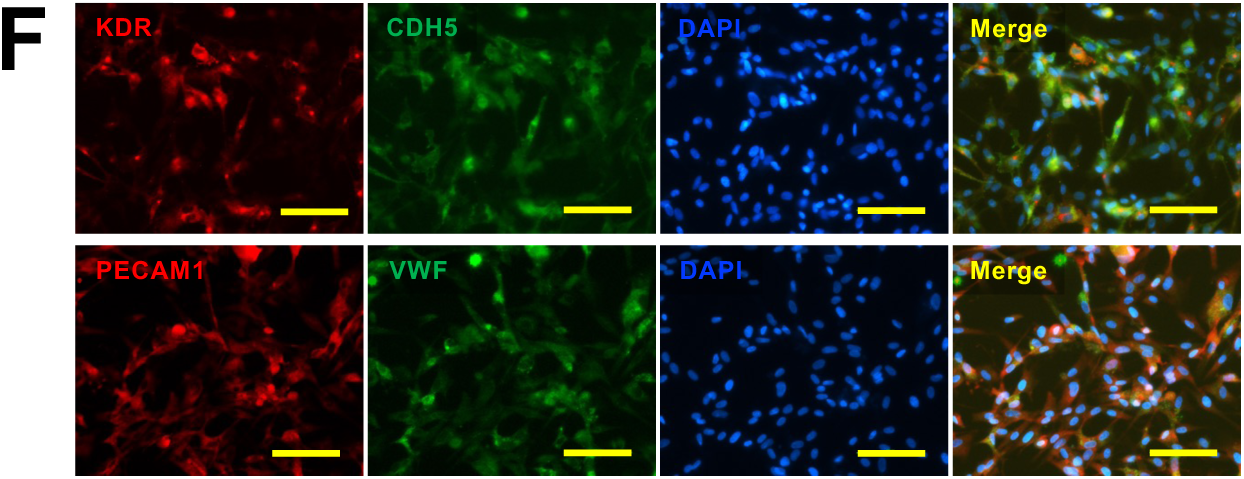
Adenoviral ETV2 rapidly reprograms human dermal fibroblasts into endothelial cells. **(A)** Schematic overview of Ad-rEC generation protocol. Human dermal fibroblasts (HDFs) are transduced with adenoviral ETV2 (MOI 70) and cultured in reprogramming medium containing ascorbic acid and SB431542 for 6 days before KDR-based FACS sorting. **(B)** Representative phase-contrast images showing morphological transformation from fibroblast to endothelial cobblestone morphology over 9 days. Scale bar, 400 μm. **(C)** Temporal expression kinetics of endothelial marker genes (*CDH5*, *KDR*, *PECAM1*) analyzed by qRT-PCR (n = 5 independent experiments). **(D)** Flow cytometric quantification of endothelial marker expression at indicated timepoints. **(E)** Comparative qRT-PCR analysis of endothelial gene expression in pre-sorted, KDR^+^, and KDR^−^ populations at day 6. Data are mean ± s.e.m. (n = 5 biological replicates). *P values determined by two-tailed unpaired t-test with Welch’s correction. **(F)** Immunofluorescence staining confirming endothelial marker expression (KDR, CDH5, PECAM1, VWF) in KDR^+^ sorted Ad-rECs. Scale bar, 100 μm.

To assess endothelial gene induction (*CDH5*, *KDR*, and *PECAM1*), we performed quantitative RT-PCR at days 2, 4, 6, and 9 post-transduction. Expression of all three endothelial genes increased throughout the culture period, with *CDH5* and *PECAM1* showing continuous upregulation over 9 days, while *KDR* expression peaked at day 6 (Fig. 1C). Flow cytometric analysis at days 4, 6, and 9 revealed that CDH5-positive cells ranged from 39% to 47%, with maximal expression at day 6. Similarly, KDR-positive cells comprised 32% to 46% of the population, also peaking at day 6. PECAM1-positive cells increased progressively from 4% to 8% over the 9-day period (Fig. 1D).

Given the peak expression of endothelial markers at day 6, we performed fluorescence-activated cell sorting (FACS) using KDR, a comprehensive endothelial-lineage marker, to isolate reprogrammed cells from Ad-ETV2-transduced HDFs (Fig. 1A). The sorted KDR^+^ population exhibited significantly elevated endothelial gene expression compared to both unsorted cells and the KDR^−^ fraction (Fig. 1E). Immunocytochemistry confirmed robust expression of multiple endothelial markers (KDR, CDH5, PECAM1, and VWF) in KDR^+^ cells (Fig. 1F). These results demonstrate that Ad-ETV2 effectively induces endothelial gene and protein expression, with KDR^+^ selection enriching for cells with endothelial characteristics.

We next evaluated whether Ad-ETV2-transduced HDFs acquired functional endothelial and proangiogenic properties. To assess endothelial functionality, we examined the cells’ ability to internalize acetylated low-density lipoprotein (Ac-LDL) and bind endothelial-specific lectins. Using CM-DiI-labeled Ac-LDL and FITC-conjugated *Bandeiraea simplicifolia* lectin 1 (BSL1) as dual markers for functional endothelial cells, we observed that approximately 40% of cells at day 9 demonstrated Ac-LDL uptake (red fluorescence), with the majority of these cells also binding BSL1 lectin (green fluorescence) (Fig. 2A). Flow cytometric quantification confirmed that 46% of cells were positive for DiI-labeled Ac-LDL (Fig. 2B).

**Figure 2.**
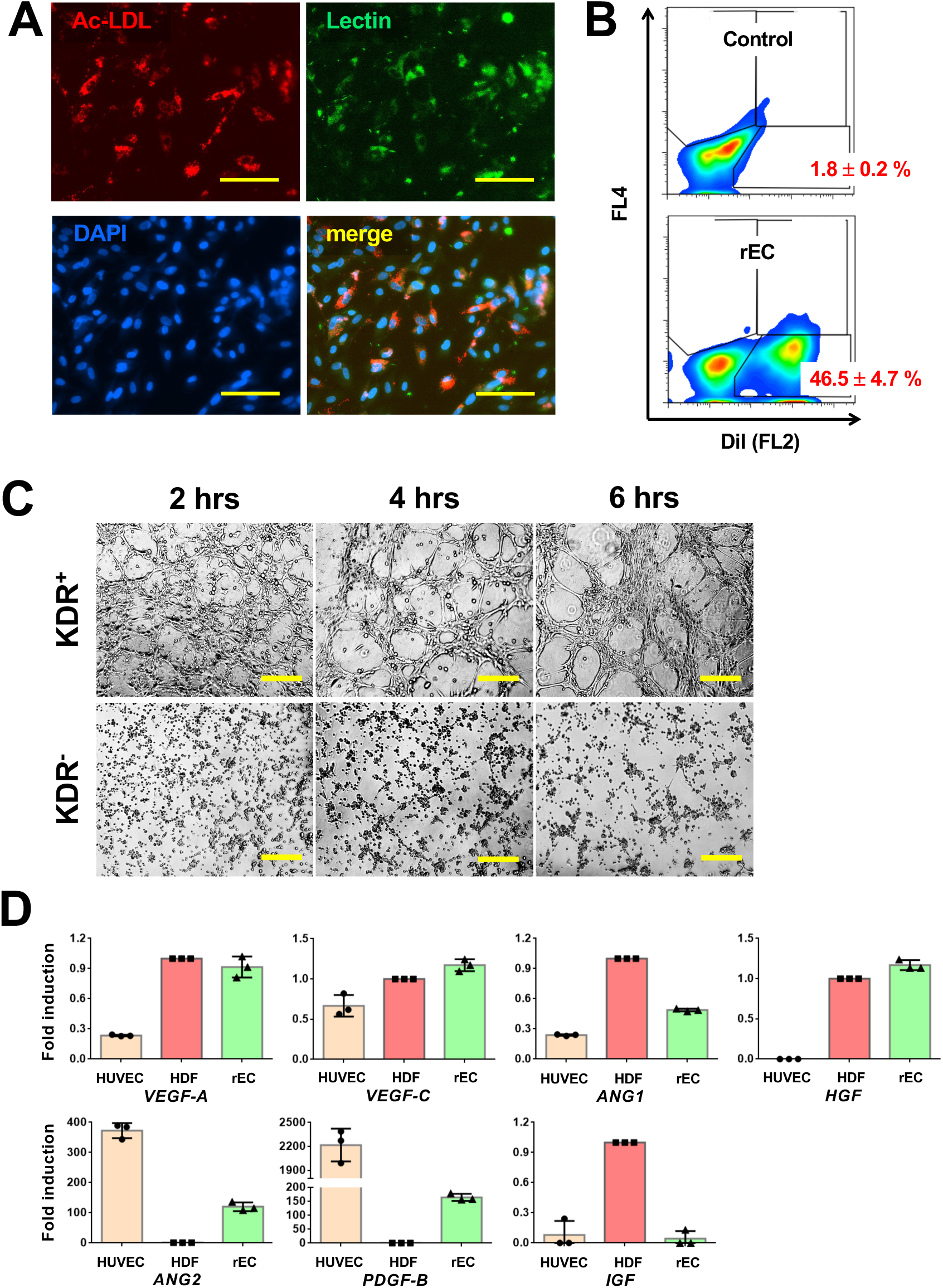
Functional characterization reveals robust endothelial and proangiogenic properties of Ad-rECs. **(A)** Dual staining for acetylated LDL uptake (CM-DiI, red) and endothelial-specific lectin binding (FITC-BSL1, green) in day 9 Ad-rECs demonstrates functional endothelial phenotype. Scale bar, 100 μm. **(B)** Flow cytometric quantification of DiI-Ac-LDL^+^ cells. **(C)** Time-course imaging of tube formation on Matrigel comparing KDR^+^ and KDR^−^ populations. Scale bar, 400 μm. **(D)** Comparative expression analysis of proangiogenic factors in Ad-rECs versus HUVECs and HDFs by qRT-PCR, revealing upregulation of therapeutic growth factors.

To evaluate angiogenic potential, we compared the tube-forming capacity of KDR^+^ and KDR^−^ populations on Matrigel. KDR^+^ cells formed robust tubular networks, while KDR^-^ cells failed to organize into tube-like structures (Fig. 2C). Analysis of proangiogenic gene expression revealed that KDR^+^ cells expressed markedly elevated levels of key angiogenic factors compared to human umbilical vein endothelial cells (HUVECs): *VEGF-A* (4-fold higher), *VEGF-C* (2-fold higher), *ANG1* (2-fold higher), and notably *HGF* (200-fold higher). Additionally, *ANG2* and *PDGF-B* expression increased more than 100-fold relative to parental HDFs (Fig. 2D). These findings demonstrate that Ad-ETV2 successfully reprograms fibroblasts into functional endothelial cells with potent proangiogenic properties, hereafter designated as Ad-rECs (adenovirally-reprogrammed endothelial cells).

### Global transcriptome analysis using RNA-sequencing

We performed RNA sequencing to comprehensively characterize Ad-rEC identity, analyzing cells at days 0 (HDFs), 1.5, 4, and 6 (including sorted KDR^+^ and KDR^−^ populations) (Fig. 3A). Principal component analysis revealed maximal transcriptional changes between days 0 and 1.5, with minimal changes between days 4 and 6 (34 upregulated, 16 downregulated genes). KDR sorting clearly separated populations, with yellow arrows indicating the reprogramming trajectory (Fig. 3A).

**Figure 3.**
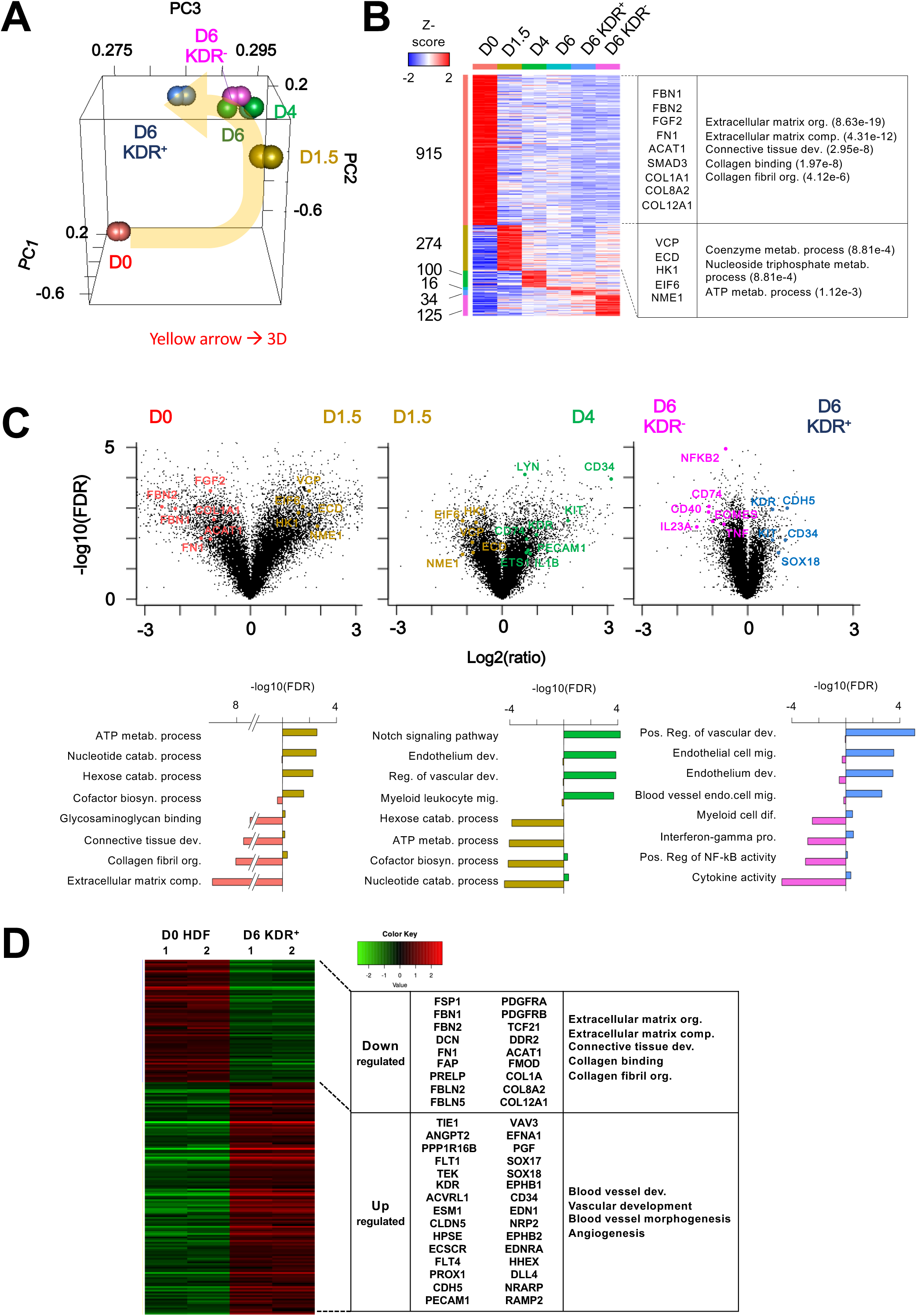
Transcriptomic profiling reveals rapid sequential reprogramming through distinct molecular phases. **(A)** Principal component analysis of RNA-seq data from reprogramming time course (days 0, 1.5, 4, 6) with biological duplicates. Yellow arrow indicates reprogramming trajectory. **(B)** Hierarchical clustering heatmap of differentially expressed genes across reprogramming timeline with representative GO terms highlighted. **(C)** Volcano plots depicting transcriptional dynamics between sequential timepoints and KDR-sorted populations. **(D)** Comparative transcriptome analysis between parental HDFs (day 0) and Ad-rECs (day 6 KDR^+^) with corresponding GO enrichment analysis demonstrating successful endothelial conversion.

Next, we analyzed differentially expressed genes (DEGs) at each time point using the Limma package pipeline^40^ and visualized in a heat map (Fig. 3B). DEG analysis at each time point revealed that the biggest transcriptional change occurred between D0 and D1.5 suggesting reprogramming can occur immediately after ETV2 overexpression. In particular, the expression of fibroblast-related genes, such as extracellular matrix and collagen related genes, were quickly suppressed at D1.5, whereas many metabolic genes were upregulated at this time point. To examine transcriptional dynamics during reprogramming, we analyzed differentially expressed genes (DEGs) between sequential timepoints: day 0 versus day 1.5, day 1.5 versus day 4, and day 6 KDR^−^ versus KDR^+^ populations. Volcano plot analysis (Fig. 3C) and Gene Ontology enrichment (Fig. 3D) revealed distinct transcriptional waves during early reprogramming. Between days 0 and 1.5, fibroblast-associated genes including *FGF2*, *FBN2*, *FBN1*, *COL1A1*, *FN1*, and *ACAT1* were significantly downregulated, while metabolic genes such as *VCP*, *EIF6*, *ECD*, *HK1*, and *NME1* were upregulated, indicating rapid metabolic reprogramming during the initial phase.

During the subsequent reprogramming phase between days 1.5 and 4, we observed a marked shift in transcriptional programs. The metabolic genes that were initially upregulated became suppressed, while genes associated with both endothelial and myeloid lineages showed increased expression. This pattern indicates that the metabolic gene activation represents a transient reprogramming phase rather than a stable cellular state. Notably, the period between days 4 and 6 showed minimal transcriptional changes, suggesting that the major reprogramming events occur within the first four days following Ad-ETV2 transduction.

Comparative analysis of KDR^+^ and KDR^−^ populations at day 6 revealed distinct lineage commitments. KDR^+^ cells demonstrated enrichment of endothelial-specific genes including *KDR*, *CDH5*, *KIT*, *CD34*, and the transcription factor *SOX18*, confirming their endothelial identity. Conversely, KDR^−^ cells exhibited elevated expression of myeloid-associated genes such as *NFKB2*, *CD74*, *CD40*, *IL23A*, *EOMES*, and *TNF*, suggesting divergent differentiation toward a myeloid-like phenotype. These distinct expression profiles demonstrate that KDR selection effectively enriches for cells with endothelial characteristics while excluding cells with alternative lineage specifications.

Comprehensive transcriptome comparison between parental fibroblasts (day 0 HDFs) and Ad-rECs (day 6 KDR^+^ cells) confirmed successful endothelial reprogramming (Fig. 3D). Fibroblast-associated genes showed consistent downregulation, while endothelial-specific genes were significantly upregulated in Ad-rECs. Gene Ontology enrichment analysis further validated this lineage conversion, revealing that Ad-rECs were significantly enriched for genes involved in vascular development and angiogenesis pathways, while showing depletion of genes encoding extracellular matrix components and collagen-related proteins characteristic of fibroblasts. These transcriptomic changes were independently validated by quantitative RT-PCR, confirming suppression of fibroblast marker genes (Supplementary Fig. S2). Collectively, these results demonstrate that Ad-ETV2 overexpression efficiently reprograms human fibroblasts into endothelial cells within six days, achieving robust induction of endothelial characteristics while simultaneously suppressing the original fibroblast identity.

### Biomaterial-enhanced therapeutic efficacy in hindlimb ischemia

Given the critical importance of cell survival for therapeutic efficacy, we first assessed the biocompatibility of PA-RGDS nanomatrix gel^38,41–43^ with Ad-rECs. Ad-rECs were pre-labeled with CM-DiI and encapsulated within PA-RGDS, then cultured under standard endothelial conditions for seven days. LIVE/DEAD staining revealed that more than 95% of encapsulated Ad-rECs remained viable throughout the culture period, demonstrating excellent biocompatibility between the nanomatrix gel and reprogrammed endothelial cells (Supplementary Fig. S3).

We next investigated the therapeutic efficacy of Ad-rECs with or without PA-RGDS encapsulation using a well-established murine hindlimb ischemia model. Ischemia was surgically induced in athymic nude mice through femoral artery ligation, followed by immediate intramuscular injection of various treatment groups: Ad-rECs alone, Ad-rECs encapsulated in PA-RGDS (Ad-rEC/PA-RGDS), HDFs transduced with control adenovirus, PA-RGDS gel alone, or PBS vehicle control. Each cellular treatment group received 4×10^5^ cells, with all cells pre-labeled with CM-DiI for subsequent histological tracking (Fig. 4A). Serial assessment of blood perfusion using laser Doppler perfusion imaging at days 0, 3, 7, 14, 21, and 28 post-surgery demonstrated significantly enhanced blood flow recovery in both Ad-rEC and Ad-rEC/PA-RGDS groups compared to all control groups (Fig. 4B,C). Notably, PA-RGDS encapsulation did not confer additional functional benefit over Ad-rECs alone during this 28-day observation period, potentially due to the relatively rapid endogenous recovery kinetics characteristic of this acute ischemia model.

**Figure 4.**
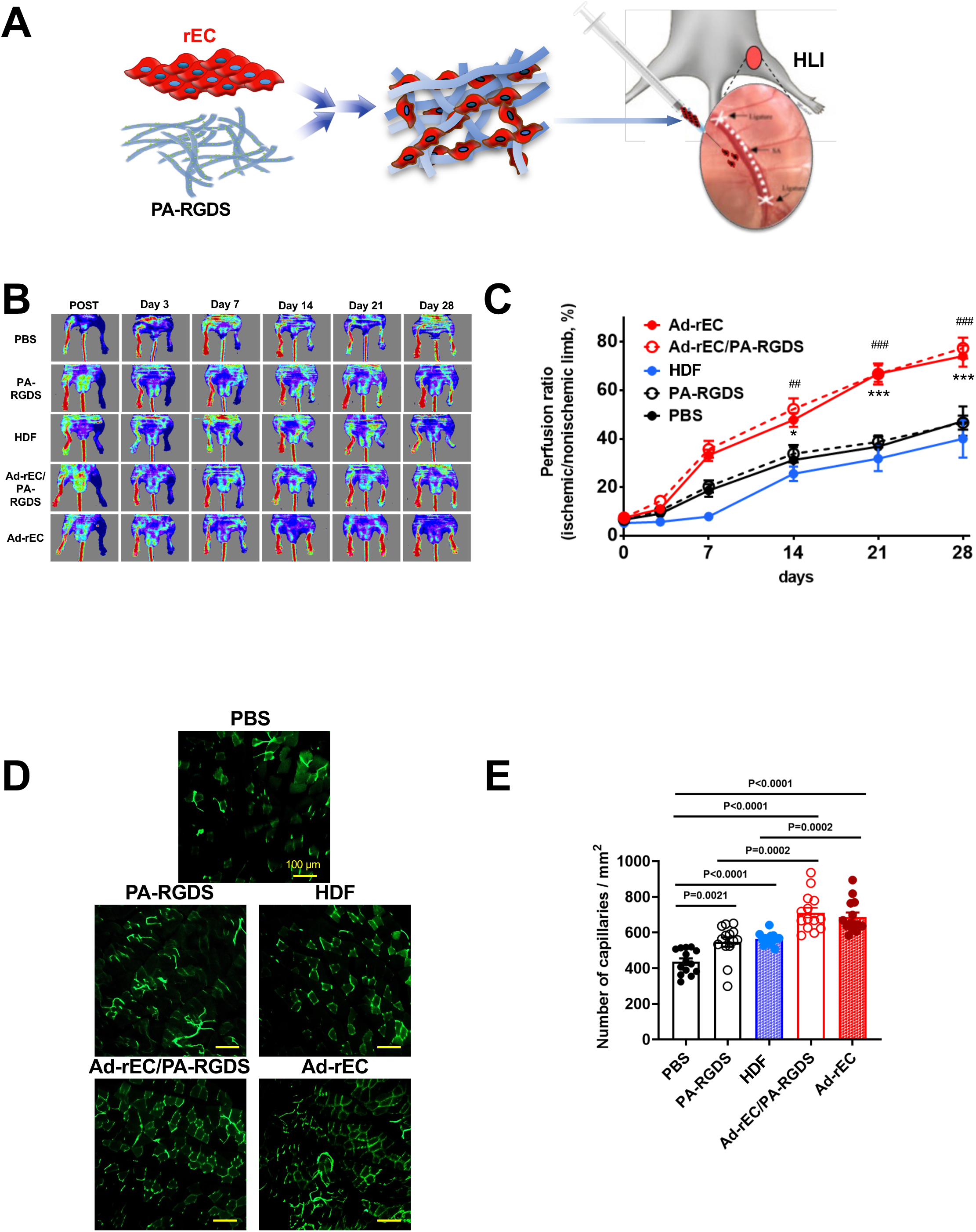
Ad-rECs promote vascular recovery and neovascularization in hindlimb ischemia. **(A)** Experimental design for therapeutic evaluation in murine hindlimb ischemia model. **(B)** Representative laser Doppler perfusion images showing blood flow recovery over 28 days. **(C)** Quantitative analysis of perfusion recovery. Data are mean ± s.e.m.; *P < 0.05, ***P < 0.0001 Ad-rEC versus controls; ^##^P < 0.001, ^###^P < 0.0001 Ad-rEC/PA-RGDS versus controls; two-way repeated measures ANOVA with Tukey’s multiple comparisons (n = 8–11 mice per group). **(D)** Representative confocal images of FITC-BSL1-perfused capillaries in ischemic muscle at day 28. Scale bar, 100 μm. **(E)** Quantification of capillary density. n = 14 fields from 5 mice per group; two-tailed unpaired t-test with Welch’s correction.

Histological evaluation of capillary density at day 28 was performed following systemic perfusion with FITC-conjugated BSL1 to label functional vasculature. Both Ad-rEC treatment groups exhibited significantly increased capillary density compared to control groups, confirming the pro-angiogenic effects of Ad-rECs in ischemic tissue (Fig. 4D,E). Interestingly, both PA-RGDS alone and HDF control groups demonstrated modestly elevated capillary density compared to PBS, suggesting some intrinsic pro-angiogenic properties of these treatments. These findings establish that Ad-rECs effectively promote vascular recovery and neovascularization in ischemic tissue, although PA-RGDS encapsulation did not enhance these short-term functional outcomes.

### Long-term engraftment and sustained vascular contribution

We assessed engraftment and vessel-forming capacity of Ad-rECs with or without PA-RGDS encapsulation through comprehensive histological examination over twelve months (4 weeks, 3, 6, and 12 months)—the longest follow-up reported for directly reprogrammed cells. Confocal microscopy revealed substantial Ad-rEC engraftment in ischemic tissue, with consistently higher retention in PA-RGDS-encapsulated versus bare Ad-rEC groups at all timepoints (Fig. 5A-H; Supplementary Fig. S4).

**Figure 5.**
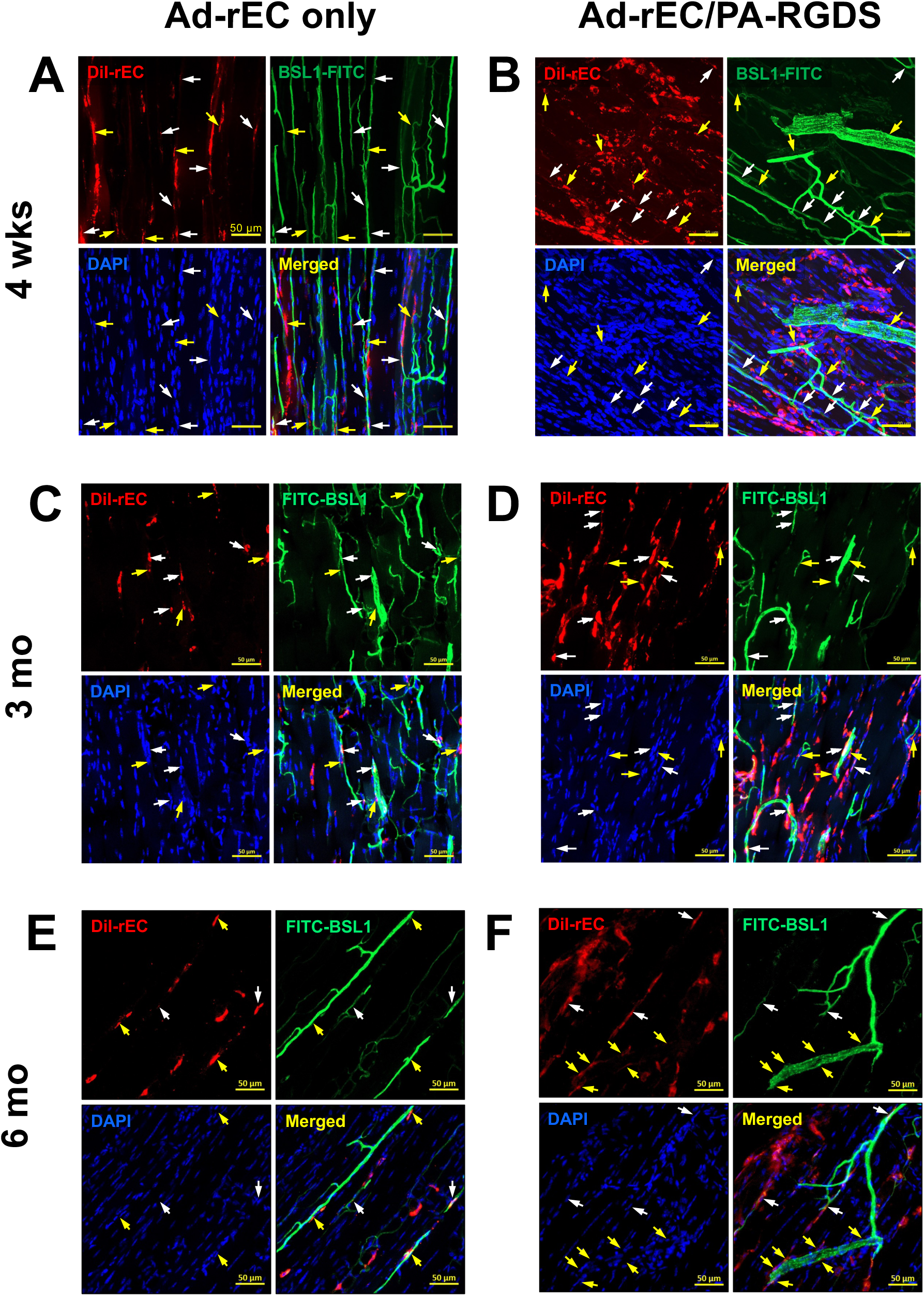
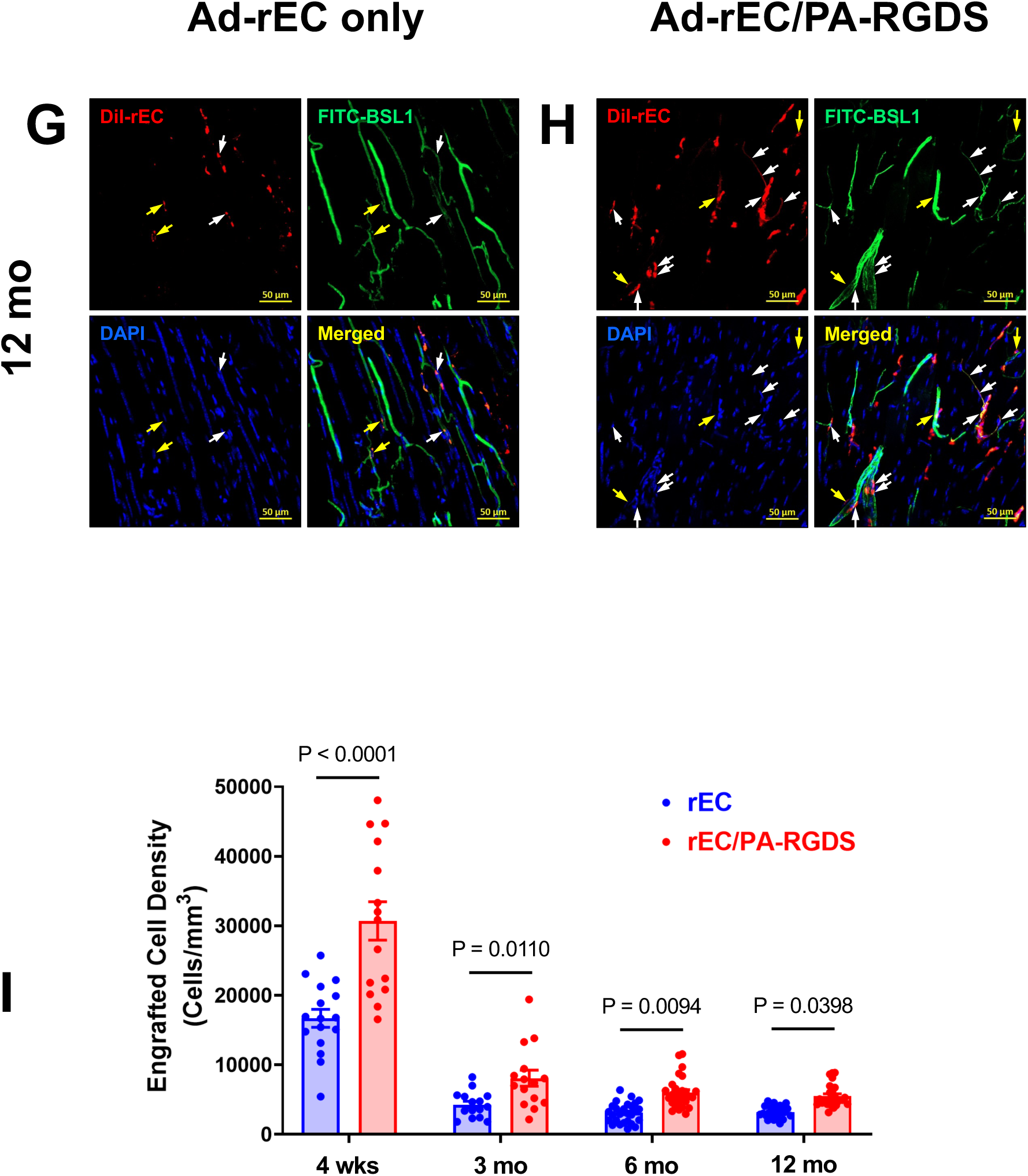
Long-term engraftment analysis demonstrates sustained vascular integration over twelve months. **(A-H)** Confocal microscopy of ischemic muscle showing CM-DiI-labeled Ad-rECs (red) and BSL1-perfused vessels (green) from mice receiving Ad-rECs alone (A, C, E, G) or Ad-rEC/PA-RGDS (B, D, F, H) at 4 weeks (A, B), 3 months (C, D), 6 months (E, F), and 12 months (G, H) post-injection. White arrows indicate vascular-incorporated Ad-rECs; yellow arrows indicate perivascular Ad-rECs. Scale bars, 50 μm. **(I)** Temporal quantification of engrafted cell density demonstrating enhanced retention with PA-RGDS encapsulation. n = 15–27 fields from 5–7 mice per group per timepoint. Mean ± s.e.m.; two-way ANOVA with Fisher’s LSD test.

While engrafted cell numbers declined over twelve months in both groups, the rate of cell loss decreased markedly over time (Fig. 5I). At three months, approximately 25% of initial cells remained; however, between three and six months, 75% of cells were retained, with similar retention between six and twelve months. This stabilization pattern occurred in both treatment groups, though absolute cell numbers remained higher with PA-RGDS encapsulation throughout the study period. These findings demonstrate that PA-RGDS nanomatrix gel significantly enhances long-term Ad-rEC retention in ischemic tissue.

Regardless of encapsulation status, Ad-rECs exhibited robust vessel-forming capacity through three distinct mechanisms. Cells migrated to perivascular regions (yellow arrows), directly incorporated into functional blood vessels as endothelial cells (white arrows), and demonstrated vessel-guiding functions (white dotted lines) (Fig. 6; Supplementary Fig. S5). These diverse vascular contributions persisted throughout the twelve-month observation period: 4 weeks (Fig. 5A,B), 3 months (Fig. 5C,D), 6 months (Fig. 5E,F), and 12 months (Fig. 5G,H) post-injection. Collectively, these data establish that Ad-rECs not only possess direct vasculogenic capacity but also contribute to vascular remodeling through perivascular support and vessel guidance functions.

**Figure 6.**
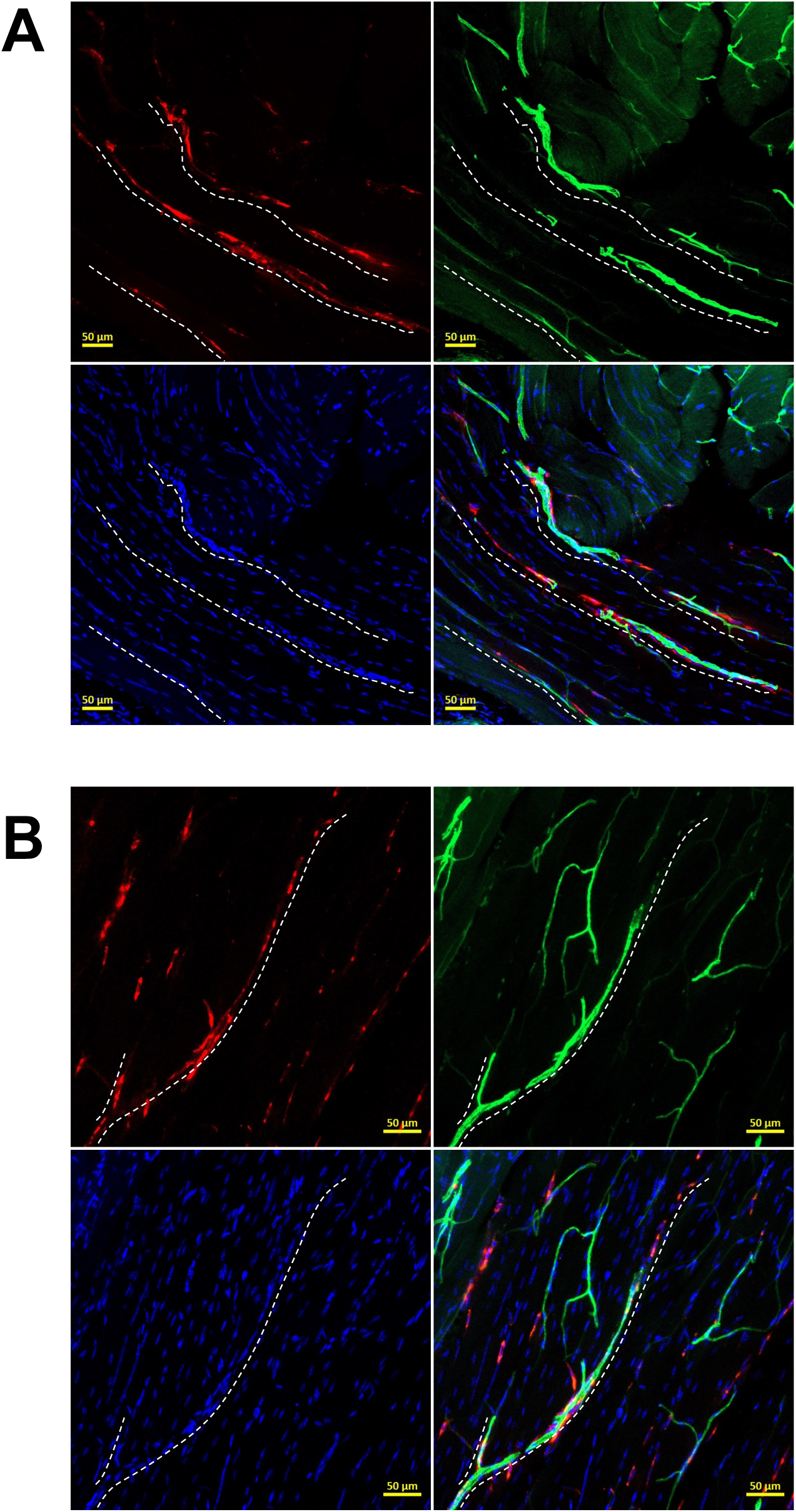
Ad-rECs contribute to neovascularization through multiple mechanisms including vessel guidance. **(A, B)** High-resolution confocal imaging at 6 months post-injection revealing three distinct mechanisms of vascular contribution in mice receiving Ad-rECs alone (A) or Ad-rEC/PA-RGDS (B). CM-DiI-labeled Ad-rECs (red) interact with BSL1-perfused vessels (green). White dotted lines highlight Ad-rECs providing structural guidance for nascent vessel formation. Scale bars, 50 μm.

## DISCUSSION

We successfully generated clinically-compatible endothelial cells through adenoviral-mediated ETV2 delivery, achieving rapid reprogramming within 48 hours with 40-50% efficiency. Global transcriptome analysis revealed sequential transcriptional waves: immediate fibroblast identity suppression, transient metabolic activation, and robust endothelial gene induction. Notably, KDR^+^ Ad-rECs expressed exceptionally high levels of proangiogenic factors, including 200-fold elevated HGF compared to HUVECs, suggesting superior therapeutic potential. Ad-rECs demonstrated potent vascular regeneration in hindlimb ischemia models, with PA-RGDS encapsulation significantly enhancing long-term engraftment—while only 25% of cells survived to three months, retention stabilized thereafter with minimal additional loss through twelve months. Ad-rECs contributed to neovascularization through three distinct mechanisms—direct vascular incorporation, perivascular support, and vessel guidance—all sustained over twelve months, representing the longest follow-up of directly reprogrammed cells to date. This non-integrating approach overcomes the primary translational barrier of genomic integration, establishing a clinically viable pathway for endothelial cell therapy in ischemic cardiovascular diseases.

Our recent study demonstrated that overexpression ETV2, a major transcription factor governing EC differentiation during development, directly reprogrammed human fibroblasts into ECs. Despite the successful generation of reprogrammed EC (rECs), lentiviral vector for ETV2 gene delivery rendered rECs clinically unsuitable. In early and recent studies of cellular reprogramming ranging from iPSCs to iCMs and rECs, lentiviral and retroviral vector have been utilized to deliver exogenous genes of transcription factors. However, these vectors induce insertional mutation on host genome by integrating exogenous genes they are carrying. Therefore, reprogrammed cells generated by lentiviral or retroviral vectors cannot be used in a clinical setting, and other methods have been sought for clinical use of reprogrammed cells. Thus, in this study, we have utilized adenovirus Ad5, which has been used for gene therapy for clinical studies to minimize the possibility of genetic integration of exogenous *ETV2* gene.

This adenoviral *ETV2* (Ad-*ETV2*) shows a robust capability to induce EC gene expression in transduced HDFs as confirmed by qRT-PCR and flow cytometry. To enrich EC lineage cell population, we sorted cells with KDR at D6 when the cell surface expression of KDR and CDH5 peaked. These KDR^+^ cells displayed significantly higher expression of endothelial genes, such as *CDH5, KDR, PECAM1, VWF,* and *NOS3*, compared to KDR^−^ cells. Moreover, transcriptome analysis of global gene expression demonstrated that the expression of vascular development and angiogenesis-related genes was highly up-regulated in KDR^+^ cells compared to HDFs, whereas the expression of fibroblast related genes such as extracellular matrix and collagen related genes was down-regulated. Also, the expression of fibroblast genes was significantly reduced in KDR^+^ cells demonstrating the loss of fibroblast characteristics accompanied by the gain of endothelial characteristics. These Ad-rECs (KDR^+^ cells) also exhibited functional characteristics of ECs such as Ac-LDL uptake and formation of tube-like structure, and expressed substantial level of proangiogenic genes. All such characteristics represent Ad-rECs as an endothelial cells.

Ad-rECs displayed therapeutic effects in an ischemic animal model. For comprehensive evaluation of therapeutic and vascularization effects of Ad-rECs, we employed a mouse hindlimb ischemia model. In a HLI, we demonstrated that injection of Ad-rECs enhanced the recovery of blood flow in the ischemic tissue. Histological analysis of Ad-rEC injected mice exhibited the increased capillary density in hindlimb compared to control mice suggesting Ad-rECs are angiogenic. This observation is also congruent with the expression of proangiogenic genes in Ad-rECs. As mentioned, cell retention is the biggest hurdle for cell therapy. Thus, we combined PA-RGDS nanomatrix gel (we encapsulated Ad-rECs in PA-RGDS nanomatrix gel) to enhance cell retention and therapeutic effects. However, we did not observe the effect of PA-RGDS on therapeutic potential of Ad-rECs for recovery of blood flow in ischemic hindlimb assessed by LDPI at 4 weeks. However, the histological analysis of ischemic tissue displayed the PA-RGDS nanomatrix gel improves the Ad-rEC retention in the hindlimb tissue. This enhanced retention of Ad-rECs with PA-RGDS was observed in the long-term follow-up over 12 months. In these ischemic tissues, the transplanted Ad-rECs were incorporated into the functional blood vessels and localized in the perivascular region suggesting that Ad-rECs contribute to neovascularization by directly taking a part of vessels as well as guiding vessel formation in the perivascular region, as we discussed in the previous study using iPSC-EC^38^. Such potent and sustained vascular regeneration effects were first shown in this study.

RNA-seq analysis of cells collected at different stages during reprogramming into Ad-rECs by *ETV2* revealed important clues for reprogramming mechanisms. First, by the comparison of DEGs between HDFs (D0) and Ad-rECs (D6 KDR^+^), and GO analysis of DEGs, we demonstrated that fibroblast-related genes were down-regulated, and EC-related and vascular development related genes were enriched in Ad-rECs. The transcriptomic analysis of global gene expression of cells collected in the course of reprogramming showed that a major change of transcriptional events occurred within 1.5 days of reprogramming after *ETV2* overexpression followed by the transcriptional induction toward endothelial lineage: during reprogramming into Ad-rEC, first, fibroblast identity is erased quickly, then metabolic genes are momentarily induced, both endothelial and myeloid gene are induced, and with presentation of KDR, cells suppressed the myeloid genes and displayed EC-like characteristics (Supplementary Fig. S6). Although these results depict the overall transcriptional landscape during the reprogramming process, it is necessary to further investigate the reprogramming events in the time course with additional experiments such as epigenetic changes.

Taken together, using adenoviral vector harboring *ETV2* (Ad-*ETV2*), we successfully generated clinically useful rECs (Ad-rECs) displaying molecular and cellular characteristics of ECs, and therapeutic effects in hindlimb ischemic mouse models. And histological data suggests that these Ad-rECs can promote neovascularization in the ischemic tissue through angiogenesis and direct contribution to vessel formation.

The rapid reprogramming kinetics observed here—with major transcriptional changes within 36 hours and functional endothelial characteristics by day 6—represents a significant advance over previous protocols. While Han et al.^15^ required 28 days using five transcription factors, and Ginsberg et al.^14^ needed 14 days with a three-factor combination, our single-factor approach achieves functional reprogramming in less than one week. This efficiency likely reflects ETV2’s role as a master regulator of endothelial development^18–25^, capable of orchestrating the entire endothelial gene program independently. The sequential transcriptional waves we identified—immediate fibroblast suppression, transient metabolic activation, and endothelial specification—provide a roadmap for understanding and potentially optimizing direct reprogramming protocols.

A particularly striking finding is the exceptional proangiogenic factor expression in Ad-rECs, with HGF levels 200-fold higher than HUVECs. This enhanced paracrine signaling capacity may explain the robust therapeutic effects at an early therapeutic effects. Recent studies by Paik et al.^44^ and Ginsberg et al.^14^ reported similar paracrine effects with reprogrammed ECs, but neither quantified factor expression to this extent nor demonstrated long-term efficacy. The elevated expression of multiple angiogenic factors (VEGF-A, VEGF-C, ANG1, ANG2, PDGF-B) suggests Ad-rECs function as “potent-secretors,” providing therapeutic benefit beyond their direct vascular contribution.

The identification of three distinct mechanisms of vascular contribution—direct incorporation, perivascular support, and vessel guidance—reveals unexpected cellular plasticity in reprogrammed cells. This multifaceted behavior resembles that of native endothelial progenitor cells and mesoangioblasts^45,46^, suggesting that Ad-rECs may recapitulate developmental vascularization programs rather than simply replacing endothelial cells. The perivascular localization particularly intrigues, as it suggests potential pericyte-like functions that could stabilize nascent vessels^47^. This finding aligns with recent work by Wakabayashi et al.^48^ showing that reprogrammed cells can exhibit multiple vascular phenotypes depending on microenvironmental cues.

Cell retention remains the Achilles’ heel of cell therapy, with most studies reporting >70% cell loss within one week^49–52^. Our observation that 75% of cells are lost by three months, but retention stabilizes thereafter, suggests a critical therapeutic window during which surviving cells establish a stable niche. The dramatic improvement in retention with PA-RGDS encapsulation— maintaining significantly higher cell numbers throughout twelve months—validates the importance of biomaterial support for clinical translation. While short-term functional outcomes (28 days) were similar between bare and encapsulated cells, the enhanced long-term retention likely translates to sustained therapeutic benefit in chronic ischemia, where extended cellular support is crucial^53,54^.

Comparison with alternative approaches highlights the advantages of our strategy. While iPSC-derived ECs offer unlimited expansion potential, they require 3-6 month for iPSC generation and 2-3 weeks for differentiation and carry inherent tumorigenicity risks^5–12^. Recent advances in mRNA-based reprogramming^55^ avoid genomic integration but require repeated administration due to transient expression. Our adenoviral approach balances safety, efficiency, and practicality—achieving rapid reprogramming with sustained expression (2-3 weeks) sufficient for cellular maturation without genomic modification.

The transcriptomic insights gained here have broader implications for cellular reprogramming. The transient metabolic activation observed during early reprogramming likely reflects the energetic demands of chromatin remodeling and transcriptional reprogramming, consistent with observations in iPSC generation^56^ and direct neuronal conversion^57^. The divergence between endothelial (KDR^+^) and myeloid (KDR^−^) programs suggests that ETV2 initially activates a broader hemato-endothelial program, with subsequent environmental cues directing final cell fate. This finding aligns with developmental biology studies showing common origins of endothelial and hematopoietic lineages^58^.

Several limitations warrant consideration. The 40-50% reprogramming efficiency, while comparable or superior to other direct reprogramming studies^14–17^, leaves room for optimization. Combining ETV2 with epigenetic modifiers or small molecules, as demonstrated in cardiac reprogramming^59^, might enhance efficiency. The mouse hindlimb ischemia model, though standard, may not fully recapitulate human chronic limb-threatening ischemia, where collateral circulation and comorbidities complicate recovery^60^. Large animal studies in porcine or non-human primate models will be essential before clinical translation.

In conclusion, we have developed a clinically tractable approach for generating therapeutic endothelial cells that addresses major translational barriers while maintaining robust efficacy. The combination of non-integrating gene delivery, rapid reprogramming, exceptional proangiogenic properties, and biomaterial-enhanced retention establishes Ad-rECs as promising candidates for treating ischemic cardiovascular diseases. These findings provide a foundation for advancing toward clinical trials while offering insights into the fundamental mechanisms of cellular reprogramming and vascular regeneration.

## METHODS

### Cell culture and maintenance

Human dermal fibroblasts (HDFs; CRL-2097, ATCC) were maintained in high-glucose DMEM (Invitrogen) supplemented with 10% fetal bovine serum at 37°C with 5% CO₂. Cells were used between passages 4–6 for all experiments. Three independent HDF batches were utilized to ensure reproducibility. Following ETV2 transduction, cells were cultured in DMEM/F12 medium supplemented with growth factors as specified in the reprogramming protocol.

### Adenoviral vector construction

Adenoviral ETV2 (Ad-ETV2) was generated using the Ad5 system (Microbix Biosystems). The ETV2 open reading frame was PCR-amplified from the FUW-tetO-ETV2 lentiviral construct^16^ and cloned into pDC315 shuttle vector. HEK293 cells were co-transfected with the ETV2-containing shuttle vector and pBHGloxΔE1,3Cre genomic vector to generate recombinant adenoviral particles. Viral stocks were amplified in HEK293 cells and titered using the Adeno-X Rapid Titer Kit (Clontech) with anti-hexon antibody detection.

### Direct endothelial reprogramming

HDFs were seeded at 2×10^4^ cells/cm^2^ and transduced with Ad-ETV2 at MOI 70, determined through preliminary optimization studies. Reprogramming medium consisted of DMEM/F12 supplemented with 10% FBS, 8 ng/ml bFGF, 50 ng/ml VEGF, 100 μM ascorbic acid, and 10 μM SB431542 (Sigma-Aldrich). Medium was replaced every 48 hours. For cell sorting, day 6 cultures were dissociated with TrypLE Express (Invitrogen) and sorted for KDR^+^ populations using fluorescence-activated cell sorting (BD FACSAria III).

### Flow cytometry

Single-cell suspensions were prepared and incubated with fluorophore-conjugated antibodies for 20 minutes at 4°C in PBS containing 2% FBS. Antibodies included: CDH5-APC (17-1449-42, eBioscience), CDH5-PE (12-1449-80, eBioscience), KDR-APC (130-093-601, Miltenyi Biotec), PECAM1-PE (130-092-653, Miltenyi Biotec), and appropriate isotype controls. Data were acquired on an Accuri C6 flow cytometer (BD Biosciences) and analyzed using FlowJo software (Tree Star) with compensation and gating based on isotype controls.

### Functional endothelial assays

For acetylated LDL uptake, cells were incubated with 10 μg/ml DiI-Ac-LDL (Biomedical Technologies) for 4 hours at 37°C, followed by 20 μg/ml FITC-BSL1 lectin (Vector Laboratories) for 1 hour. For tube formation, 2×10^5^ cells were seeded on growth factor-reduced Matrigel (BD Biosciences) in EGM-2 medium and imaged at 2, 4, and 6 hours.

### Immunofluorescence microscopy

Cells were fixed with 4% paraformaldehyde, permeabilized with 0.1% Triton X-100, and blocked with 5% normal goat serum. Primary antibodies included: VWF (1:100; SC-8068, Santa Cruz), KDR (1:100; 2479, Cell Signaling), CDH5 (1:100; 14-1449, eBioscience), and PECAM1 (1:100; CBL-468, Millipore). Alexa Fluor-conjugated secondary antibodies (Invitrogen) were used for detection. Images were acquired using a Zeiss LSM 510 Meta confocal microscope.

### Gene expression analysis

RNA was extracted using RNeasy Mini Kit (Qiagen) and reverse-transcribed with TaqMan Reverse Transcription Reagents (Applied Biosystems). Quantitative PCR was performed using SYBR Green Supermix (Bio-Rad) on a 7500 Fast Real-Time PCR System (Applied Biosystems). Expression was normalized to GAPDH using the 2^−ΔΔCt^ method. Primer sequences are provided in Supplementary Table 1.

### RNA sequencing and bioinformatics

Total RNA from biological duplicates at days 0, 1.5, 4, 6 (unsorted), and day 6 KDR^+^/KDR^−^ populations underwent library preparation using TruSeq Stranded mRNA Sample Prep Kit (Illumina). Sequencing was performed on NovaSeq 6000 (150 bp paired-end reads). Reads were aligned to hg19 using TopHat2 (v2.1.1)^62^. and expression quantified with Cufflinks (v2.2.1)^63^. Differentially expressed genes were identified using 2-fold change and adjusted P < 0.05 criteria. Gene Ontology enrichment was analyzed using GOstats (v2.24.0)^64^ with Benjamini-Hochberg correction. Additional analyses utilized iDEP (v0.94)^65^. Data are deposited in GEO (accession GSE276674).

### Hindlimb ischemia model

All animal procedures were approved by [Institution] IACUC. Male athymic nude mice (12–16 weeks, Charles River) underwent unilateral femoral artery ligation with cauterization of collateral branches. Immediately post-surgery, 4×10^5^ CM-DiI-labeled cells in 100 μl PBS or encapsulated in PA-RGDS nanomatrix gel were injected intramuscularly at three sites in the ischemic gastrocnemius muscle. Treatment groups included: PBS (n=9), HDFs transduced with control adenovirus (n=9), PA-RGDS alone (n=8), Ad-rECs (n=9), or Ad-rEC/PA-RGDS (n=11). Animals were randomized using random.org and investigators were blinded to treatment groups during analysis.

### Blood perfusion assessment

Laser Doppler perfusion imaging (LDPI; Moor Instruments) was performed pre-surgery, immediately post-surgery, and at days 3, 7, 14, 21, and 28. Perfusion ratios were calculated as ischemic/contralateral limb values. Animals with >20% residual flow immediately post-surgery were excluded from analysis.

### Histological analysis

For terminal vascular labeling, mice received intracardiac injection of 100 μg FITC-BSL1 in 100 μl PBS before euthanasia. Gastrocnemius muscles were harvested, fixed in 4% paraformaldehyde, cryoprotected in 30% sucrose, and sectioned at 20 μm thickness. Capillary density was quantified from ≥5 randomly selected fields per animal using confocal microscopy (Zeiss LSM 510 Meta). Engrafted cell density was calculated from CM-DiI^+^ cells in 10 random high-power fields per section.

### PA-RGDS nanomatrix gel preparation

PA-RGDS was synthesized and characterized as previously described^38^. Briefly, palmitic acid was conjugated to GTAGLIGQ-RGDS peptide sequence via alkylation. The amphiphile was dissolved in sterile water at 1% (w/v) and induced to gel by addition of cell suspension in medium, achieving physiological pH and ionic strength. Biocompatibility was assessed using LIVE/DEAD staining (Invitrogen) after 7 days of culture.

### Statistical analysis

Sample sizes were determined using power analysis (α=0.05, β=0.90) based on preliminary data. Data normality was assessed using Shapiro-Wilk test. For two-group comparisons, unpaired two-tailed t-tests with Welch’s correction were used. Multiple group comparisons employed two-way repeated measures ANOVA with Tukey’s post-hoc test. The exclusion criterion was set at 1.5× s.e.m. from the mean; no data met this criterion. All experiments were performed with ≥3 biological replicates. Data are presented as mean ± s.e.m. Statistical analyses were performed using GraphPad Prism 9. P < 0.05 was considered significant.

### Data availability

RNA-seq data are available at GEO (GSE276674). All other data supporting the findings are available from the corresponding author upon reasonable request.

## Non-standard Abbreviation and Acronyms

EC: endothelial cell
rEC: reprogrammed endothelial cell
Ad: adenoviral
PA: peptide amphiphile
RGDS: Arg-Gly-Asp-Ser
iPSC: induced pluripotent stem cell
HLI: hindlimb ischemia

---

**Table I.**
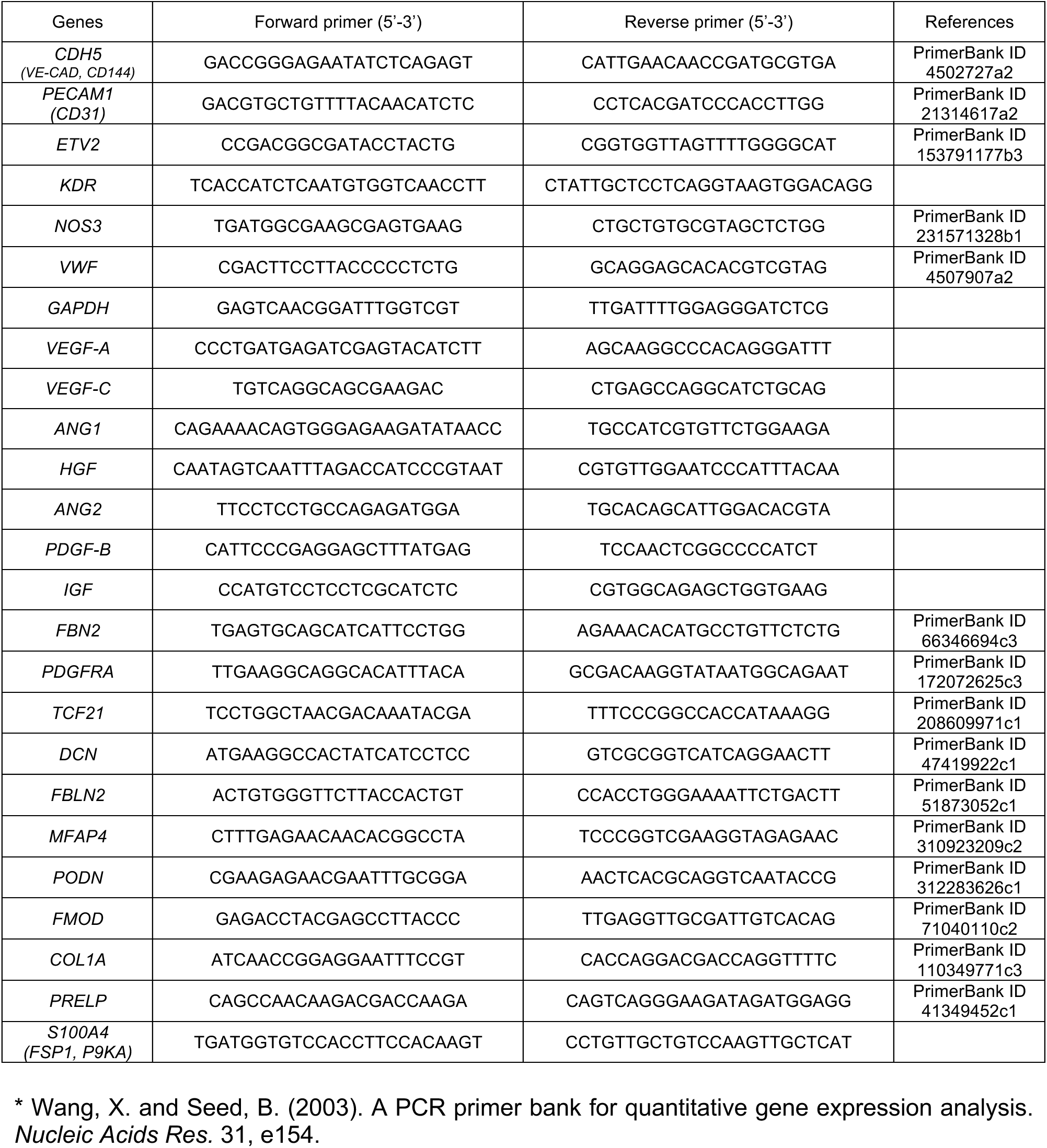
The sequences of primers used for qRT-PCR analysis, Related to Experimental Procedures. Some of the listed primers were designed according to PrimerBank (http://pga.mgh.harvard.edu/primerbank)* database.

## REFERENCES

1. Tsao CW, Aday AW, Almarzooq ZI, Anderson CAM, Arora P, Avery CL, Baker-Smith CM, Beaton AZ, Boehme AK, Buxton AE, Commodore-Mensah Y, Elkind MSV, Evenson KR, Eze-Nliam C, Fugar S, Generoso G, Heard DG, Hiremath S, Ho JE, Kalani R, Kazi DS, Ko D, Levine DA, Liu J, Ma J, Magnani JW, Michos ED, Mussolino ME, Navaneethan SD, Parikh NI, Poudel R, Rezk-Hanna M, Roth GA, Shah NS, St-Onge MP, Thacker EL, Virani SS, Voeks JH, Wang NY, Wong ND, Wong SS, Yaffe K, Martin SS, American Heart Association Council on E, Prevention Statistics C and Stroke Statistics S. Heart Disease and Stroke Statistics-2023 Update: A Report From the American Heart Association. Circulation. 2023;147:e93–e621.

2. Wong ND. Epidemiological studies of CHD and the evolution of preventive cardiology. Nature reviews Cardiology. 2014;11:276–89.

3. Losordo DW and Dimmeler S. Therapeutic angiogenesis and vasculogenesis for ischemic disease: part II: cell-based therapies. Circulation. 2004;109:2692–7.

4. Losordo DW and Dimmeler S. Therapeutic angiogenesis and vasculogenesis for ischemic disease. Part I: angiogenic cytokines. Circulation. 2004;109:2487–91.

5. Cohen DE and Melton D. Turning straw into gold: directing cell fate for regenerative medicine. Nat Rev Genet. 2011;12:243–52.

6. Knoepfler PS. Deconstructing stem cell tumorigenicity: a roadmap to safe regenerative medicine. Stem Cells. 2009;27:1050–6.

7. Yamanaka S. A fresh look at iPS cells. Cell. 2009;137:13–7.

8. Cho SW, Moon SH, Lee SH, Kang SW, Kim J, Lim JM, Kim HS, Kim BS and Chung HM. Improvement of postnatal neovascularization by human embryonic stem cell derived endothelial-like cell transplantation in a mouse model of hindlimb ischemia. Circulation. 2007;116:2409–19.

9. Li Z, Hu S, Ghosh Z, Han Z and Wu JC. Functional characterization and expression profiling of human induced pluripotent stem cell- and embryonic stem cell-derived endothelial cells. Stem cells and development. 2011;20:1701–10.

10. Levenberg S, Golub JS, Amit M, Itskovitz-Eldor J and Langer R. Endothelial cells derived from human embryonic stem cells. Proceedings of the National Academy of Sciences of the United States of America. 2002;99:4391–6.

11. Choi KD, Yu J, Smuga-Otto K, Salvagiotto G, Rehrauer W, Vodyanik M, Thomson J and Slukvin I. Hematopoietic and endothelial differentiation of human induced pluripotent stem cells. Stem Cells. 2009;27:559–67.

12. Wang ZZ, Au P, Chen T, Shao Y, Daheron LM, Bai H, Arzigian M, Fukumura D, Jain RK and Scadden DT. Endothelial cells derived from human embryonic stem cells form durable blood vessels in vivo. Nat Biotechnol. 2007;25:317–8.

13. Jeong JO, Han JW, Kim JM, Cho HJ, Park C, Lee N, Kim DW and Yoon YS. Malignant tumor formation after transplantation of short-term cultured bone marrow mesenchymal stem cells in experimental myocardial infarction and diabetic neuropathy. Circ Res. 2011;108:1340–7.

14. Ginsberg M, James D, Ding BS, Nolan D, Geng F, Butler JM, Schachterle W, Pulijaal VR, Mathew S, Chasen ST, Xiang J, Rosenwaks Z, Shido K, Elemento O, Rabbany SY and Rafii S. Efficient direct reprogramming of mature amniotic cells into endothelial cells by ETS factors and TGFbeta suppression. Cell. 2012;151:559–75.

15. Han JK, Shin Y, Sohn MH, Choi SB, Shin D, You Y, Shin JY, Seo JS and Kim HS. Direct conversion of adult human fibroblasts into functional endothelial cells using defined factors. Biomaterials. 2021;272:120781.

16. Lee S, Park C, Han JW, Kim JY, Cho K, Kim EJ, Kim S, Lee SJ, Oh SY, Tanaka Y, Park IH, An HJ, Shin CM, Sharma S and Yoon YS. Direct Reprogramming of Human Dermal Fibroblasts Into Endothelial Cells Using ER71/ETV2. Circ Res. 2017;120:848–861.

17. Morita R, Suzuki M, Kasahara H, Shimizu N, Shichita T, Sekiya T, Kimura A, Sasaki K, Yasukawa H and Yoshimura A. ETS transcription factor ETV2 directly converts human fibroblasts into functional endothelial cells. Proceedings of the National Academy of Sciences of the United States of America. 2015;112:160–5.

18. De Haro L and Janknecht R. Functional analysis of the transcription factor ER71 and its activation of the matrix metalloproteinase-1 promoter. Nucleic Acids Res. 2002;30:2972–9.

19. De Haro L and Janknecht R. Cloning of the murine ER71 gene (Etsrp71) and initial characterization of its promoter. Genomics. 2005;85:493–502.

20. Ferdous A, Caprioli A, Iacovino M, Martin CM, Morris J, Richardson JA, Latif S, Hammer RE, Harvey RP, Olson EN, Kyba M and Garry DJ. Nkx2-5 transactivates the Ets-related protein 71 gene and specifies an endothelial/endocardial fate in the developing embryo. Proceedings of the National Academy of Sciences of the United States of America. 2009;106:814–9.

21. Kataoka H, Hayashi M, Nakagawa R, Tanaka Y, Izumi N, Nishikawa S, Jakt ML and Tarui H. Etv2/ER71 induces vascular mesoderm from Flk1+PDGFRalpha+ primitive mesoderm. Blood. 2011;118:6975–86.

22. Lee D, Park C, Lee H, Lugus JJ, Kim SH, Arentson E, Chung YS, Gomez G, Kyba M, Lin S, Janknecht R, Lim DS and Choi K. ER71 acts downstream of BMP, Notch, and Wnt signaling in blood and vessel progenitor specification. Cell stem cell. 2008;2:497–507.

23. Liu F, Kang I, Park C, Chang LW, Wang W, Lee D, Lim DS, Vittet D, Nerbonne JM and Choi K. ER71 specifies Flk-1+ hemangiogenic mesoderm by inhibiting cardiac mesoderm and Wnt signaling. Blood. 2012;119:3295–305.

24. De Val S, Chi NC, Meadows SM, Minovitsky S, Anderson JP, Harris IS, Ehlers ML, Agarwal P, Visel A, Xu SM, Pennacchio LA, Dubchak I, Krieg PA, Stainier DY and Black BL. Combinatorial regulation of endothelial gene expression by ets and forkhead transcription factors. Cell. 2008;135:1053–64.

25. Lee D, Kim T and Lim DS. The Er71 is an important regulator of hematopoietic stem cells in adult mice. Stem Cells. 2011;29:539–48.

26. Cho S, Aakash P, Lee S and Yoon YS. Endothelial cell direct reprogramming: Past, present, and future. J Mol Cell Cardiol. 2023;180:22–32.

27. Jung C, Oh JE, Lee S and Yoon YS. Generation and Application of Directly Reprogrammed Endothelial Cells. Korean Circ J. 2022;52:643–658.

28. Hoffmann J, Glassford AJ, Doyle TC, Robbins RC, Schrepfer S and Pelletier MP. Angiogenic effects despite limited cell survival of bone marrow-derived mesenchymal stem cells under ischemia. The Thoracic and cardiovascular surgeon. 2010;58:136–42.

29. Huang NF, Niiyama H, Peter C, De A, Natkunam Y, Fleissner F, Li Z, Rollins MD, Wu JC, Gambhir SS and Cooke JP. Embryonic stem cell-derived endothelial cells engraft into the ischemic hindlimb and restore perfusion. Arteriosclerosis, thrombosis, and vascular biology. 2010;30:984–91.

30. Huang NF, Okogbaa J, Babakhanyan A and Cooke JP. Bioluminescence imaging of stem cell-based therapeutics for vascular regeneration. Theranostics. 2012;2:346–54.

31. Laurila JP, Laatikainen L, Castellone MD, Trivedi P, Heikkila J, Hinkkanen A, Hematti P and Laukkanen MO. Human embryonic stem cell-derived mesenchymal stromal cell transplantation in a rat hind limb injury model. Cytotherapy. 2009;11:726–37.

32. Rufaihah AJ, Huang NF, Jame S, Lee JC, Nguyen HN, Byers B, De A, Okogbaa J, Rollins M, Reijo-Pera R, Gambhir SS and Cooke JP. Endothelial cells derived from human iPSCS increase capillary density and improve perfusion in a mouse model of peripheral arterial disease. Arteriosclerosis, thrombosis, and vascular biology. 2011;31:e72–9.

33. Wollert KC and Drexler H. Clinical applications of stem cells for the heart. Circ Res. 2005;96:151–63.

34. Menasche P. Skeletal myoblasts as a therapeutic agent. Progress in cardiovascular diseases. 2007;50:7–17.

35. Hofmann M, Wollert KC, Meyer GP, Menke A, Arseniev L, Hertenstein B, Ganser A, Knapp WH and Drexler H. Monitoring of bone marrow cell homing into the infarcted human myocardium. Circulation. 2005;111:2198–202.

36. Qian H, Yang Y, Huang J, Gao R, Dou K, Yang G, Li J, Shen R, He Z, Lu M and Zhao S. Intracoronary delivery of autologous bone marrow mononuclear cells radiolabeled by 18F-fluoro-deoxy-glucose: tissue distribution and impact on post-infarct swine hearts. J Cell Biochem. 2007;102:64–74.

37. Ban K, Park H-J, Kim S, Cho K-W, Hwang JW, Cha HJ, Andukuri A, Jun H-W and Yoon Y-s. Engineered Cell Therapy With Embryonic Stem Cell-Derived Cardiomyocytes Encapsulated in Injectable Nanomatrix Gel Enhanced Engraftment and Promoted Cardiac Repair in Experimental Myocardial Infarction. Circulation. 2013;128:A18111.

38. Lee SJ, Sohn YD, Andukuri A, Kim S, Byun J, Han JW, Park IH, Jun HW and Yoon YS. Enhanced Therapeutic and Long-Term Dynamic Vascularization Effects of Human Pluripotent Stem Cell-Derived Endothelial Cells Encapsulated in a Nanomatrix Gel. Circulation. 2017;136:1939–1954.

39. Wold WS and Toth K. Adenovirus vectors for gene therapy, vaccination and cancer gene therapy. Curr Gene Ther. 2013;13:421–33.

40. Ritchie ME, Phipson B, Wu D, Hu Y, Law CW, Shi W and Smyth GK. limma powers differential expression analyses for RNA-sequencing and microarray studies. Nucleic Acids Res. 2015;43:e47.

41. Anderson JM, Andukuri A, Lim DJ and Jun HW. Modulating the gelation properties of self-assembling peptide amphiphiles. ACS Nano. 2009;3:3447–54.

42. Anderson JM, Patterson JL, Vines JB, Javed A, Gilbert SR and Jun HW. Biphasic peptide amphiphile nanomatrix embedded with hydroxyapatite nanoparticles for stimulated osteoinductive response. ACS Nano. 2011;5:9463–79.

43. Ban K, Park HJ, Kim S, Andukuri A, Cho KW, Hwang JW, Cha HJ, Kim SY, Kim WS, Jun HW and Yoon YS. Cell therapy with embryonic stem cell-derived cardiomyocytes encapsulated in injectable nanomatrix gel enhances cell engraftment and promotes cardiac repair. ACS Nano. 2014;8:10815–25.

44. Paik DT, Tian L, Lee J, Sayed N, Chen IY, Rhee S, Rhee JW, Kim Y, Wirka RC, Buikema JW, Wu SM, Red-Horse K, Quertermous T and Wu JC. Large-Scale Single-Cell RNA-Seq Reveals Molecular Signatures of Heterogeneous Populations of Human Induced Pluripotent Stem Cell-Derived Endothelial Cells. Circ Res. 2018;123:443–450.

45. Dellavalle A, Sampaolesi M, Tonlorenzi R, Tagliafico E, Sacchetti B, Perani L, Innocenzi A, Galvez BG, Messina G, Morosetti R, Li S, Belicchi M, Peretti G, Chamberlain JS, Wright WE, Torrente Y, Ferrari S, Bianco P and Cossu G. Pericytes of human skeletal muscle are myogenic precursors distinct from satellite cells. Nat Cell Biol. 2007;9:255–67.

46. Medina RJ, Barber CL, Sabatier F, Dignat-George F, Melero-Martin JM, Khosrotehrani K, Ohneda O, Randi AM, Chan JKY, Yamaguchi T, Van Hinsbergh VWM, Yoder MC and Stitt AW. Endothelial Progenitors: A Consensus Statement on Nomenclature. Stem Cells Transl Med. 2017;6:1316–1320.

47. Armulik A, Genove G and Betsholtz C. Pericytes: developmental, physiological, and pathological perspectives, problems, and promises. Dev Cell. 2011;21:193–215.

48. Wakabayashi T, Naito H, Suehiro JI, Lin Y, Kawaji H, Iba T, Kouno T, Ishikawa-Kato S, Furuno M, Takara K, Muramatsu F, Weizhen J, Kidoya H, Ishihara K, Hayashizaki Y, Nishida K, Yoder MC and Takakura N. CD157 Marks Tissue-Resident Endothelial Stem Cells with Homeostatic and Regenerative Properties. Cell stem cell. 2018;22:384–397 e6.

49. Hou D, Youssef EA, Brinton TJ, Zhang P, Rogers P, Price ET, Yeung AC, Johnstone BH, Yock PG and March KL. Radiolabeled cell distribution after intramyocardial, intracoronary, and interstitial retrograde coronary venous delivery: implications for current clinical trials. Circulation. 2005;112:I150–6.

50. Teng CJ, Luo J, Chiu RC and Shum-Tim D. Massive mechanical loss of microspheres with direct intramyocardial injection in the beating heart: implications for cellular cardiomyoplasty. The Journal of thoracic and cardiovascular surgery. 2006;132:628–32.

51. Terrovitis J, Lautamaki R, Bonios M, Fox J, Engles JM, Yu J, Leppo MK, Pomper MG, Wahl RL, Seidel J, Tsui BM, Bengel FM, Abraham MR and Marban E. Noninvasive quantification and optimization of acute cell retention by in vivo positron emission tomography after intramyocardial cardiac-derived stem cell delivery. J Am Coll Cardiol. 2009;54:1619–26.

52. Zhang M, Mal N, Kiedrowski M, Chacko M, Askari AT, Popovic ZB, Koc ON and Penn MS. SDF-1 expression by mesenchymal stem cells results in trophic support of cardiac myocytes after myocardial infarction. FASEB J. 2007;21:3197–207.

53. Cooke JP and Losordo DW. Modulating the vascular response to limb ischemia: angiogenic and cell therapies. Circ Res. 2015;116:1561–78.

54. Qadura M, Terenzi DC, Verma S, Al-Omran M and Hess DA. Concise Review: Cell Therapy for Critical Limb Ischemia: An Integrated Review of Preclinical and Clinical Studies. Stem Cells. 2018;36:161–171.

55. Kaur K, Hadas Y, Kurian AA, Zak MM, Yoo J, Mahmood A, Girard H, Komargodski R, Io T, Santini MP, Sultana N, Sharkar MTK, Magadum A, Fargnoli A, Yoon S, Chepurko E, Chepurko V, Eliyahu E, Pinto D, Lebeche D, Kovacic JC, Hajjar RJ, Rafii S and Zangi L. Direct reprogramming induces vascular regeneration post muscle ischemic injury. Mol Ther. 2021;29:3042–3058.

56. Panopoulos AD, Yanes O, Ruiz S, Kida YS, Diep D, Tautenhahn R, Herrerias A, Batchelder EM, Plongthongkum N, Lutz M, Berggren WT, Zhang K, Evans RM, Siuzdak G and Izpisua Belmonte JC. The metabolome of induced pluripotent stem cells reveals metabolic changes occurring in somatic cell reprogramming. Cell Res. 2012;22:168–77.

57. Gascon S, Masserdotti G, Russo GL and Gotz M. Direct Neuronal Reprogramming: Achievements, Hurdles, and New Roads to Success. Cell stem cell. 2017;21:18–34.

58. Ditadi A, Sturgeon CM, Tober J, Awong G, Kennedy M, Yzaguirre AD, Azzola L, Ng ES, Stanley EG, French DL, Cheng X, Gadue P, Speck NA, Elefanty AG and Keller G. Human definitive haemogenic endothelium and arterial vascular endothelium represent distinct lineages. Nat Cell Biol. 2015;17:580–91.

59. Cao N, Huang Y, Zheng J, Spencer CI, Zhang Y, Fu JD, Nie B, Xie M, Zhang M, Wang H, Ma T, Xu T, Shi G, Srivastava D and Ding S. Conversion of human fibroblasts into functional cardiomyocytes by small molecules. Science. 2016;352:1216–20.

60. Padgett ME, McCord TJ, McClung JM and Kontos CD. Methods for Acute and Subacute Murine Hindlimb Ischemia. J Vis Exp. 2016.

61. Kim H, Cho HJ, Kim SW, Liu B, Choi YJ, Lee J, Sohn YD, Lee MY, Houge MA and Yoon YS. CD31+ cells represent highly angiogenic and vasculogenic cells in bone marrow: novel role of nonendothelial CD31+ cells in neovascularization and their therapeutic effects on ischemic vascular disease. Circ Res. 2010;107:602–14.

62. Trapnell C, Pachter L and Salzberg SL. TopHat: discovering splice junctions with RNA-Seq. Bioinformatics. 2009;25:1105–11.

63. Trapnell C, Williams BA, Pertea G, Mortazavi A, Kwan G, van Baren MJ, Salzberg SL, Wold BJ and Pachter L. Transcript assembly and quantification by RNA-Seq reveals unannotated transcripts and isoform switching during cell differentiation. Nat Biotechnol. 2010;28:511–5.

64. Falcon S and Gentleman R. Using GOstats to test gene lists for GO term association. Bioinformatics. 2007;23:257–8.

65. Ge SX, Son EW and Yao R. iDEP: an integrated web application for differential expression and pathway analysis of RNA-Seq data. BMC Bioinformatics. 2018;19:534.

