## Supplementary Figures for "Non-integrating direct reprogramming generates therapeutic endothelial cells with sustained vascular regeneration capacity enhanced by nanomatrix delivery"

### SUPPLEMENT DATA

#### SUPPLEMENTARY FIGURE LEGEND

**Supplementary Figure S1. Construction and validation of adenoviral ETV2 vector.** Schematic diagram illustrating Ad-ETV2 generation using the Microbix Biosystems Ad5 adenovirus system, including cloning strategy and vector elements.

**Supplementary Figure S2. Quantification of fibroblast marker suppression in Ad-rECs.** qRT-PCR analysis demonstrating significant downregulation of fibroblast-specific genes in $KDR^+$ Ad-rECs compared to parental HDFs at day 6. Data are mean ± s.e.m. (n = 5 biological replicates). *P values by two-tailed unpaired t-test with Welch's correction.

**Supplementary Figure S3. PA-RGDS nanomatrix gel demonstrates excellent biocompatibility with Ad-rECs. (A)** Brightfield and fluorescence imaging of DiI-labeled Ad-rECs encapsulated in PA-RGDS. Scale bar, 200 μm. **(B)** Live/dead staining after 7 days of culture showing >95% viability. Scale bar, 50 μm.

**Supplementary Figure S4. Extended imaging demonstrates consistent vascular contribution patterns across all timepoints. (A-D)** Additional confocal images from 4 weeks (**A**), 3 months (**B**), 6 months (**B**), and 12 months (**D**) post-injection showing CM-DiI-labeled Ad-rECs (red) and BSL1-perfused vessels (green). White arrows, vascular-incorporated cells; yellow arrows, perivascular cells. Scale bars, 50 μm.

**Supplementary Figure S5. Three-dimensional reconstruction reveals spatial organization of Ad-rEC vascular integration.** 3D-rendered confocal z-stacks from 6-month specimens showing spatial relationships between CM-DiI-labeled Ad-rECs (red) and BSL1-perfused vasculature (green) in mice receiving Ad-rECs alone or with PA-RGDS encapsulation.

**Supplementary Figure S6. Model summarizing transcriptional dynamics during ETV2-mediated endothelial reprogramming.** Schematic representation of sequential molecular phases identified by RNA-seq analysis, illustrating the progression from fibroblast suppression through metabolic activation to endothelial specification. Representative phase-contrast images from Figure 1 illustrate morphological changes. Scale bar, 400 μm.
